# Task matters: individual MEG signatures from naturalistic and neurophysiological brain states

**DOI:** 10.1101/2022.08.25.505232

**Authors:** Nigel Colenbier, Ekansh Sareen, Tamara del-Águila Puntas, Alessandra Griffa, Giovanni Pellegrino, Dante Mantini, Daniele Marinazzo, Giorgio Arcara, Enrico Amico

## Abstract

The discovery that human brain connectivity data can be used as a “fingerprint” to identify a given individual from a population, has become a burgeoning research area in the neuroscience field. Recent studies have identified the possibility to extract these brain signatures from the temporal rich dynamics of resting-state magnetoencephalography (MEG) recordings. However, to what extent MEG signatures constitute a marker of human identifiability when engaged in task-related behavior remains an open question. Here, using MEG data from naturalistic and neurophysiological tasks, we show that identification improves in tasks relative to resting-state, providing compelling evidence for a task dependent axis of MEG signatures. Notably, improvements in identifiability were more prominent in strictly controlled tasks. Lastly, the brain regions contributing most towards individual identification were also modified when engaged in task activities. We hope that this investigation advances our understanding of the driving factors behind brain identification from MEG signals.

## Introduction

The patterns of the human fingertip ridges have been established as being a “signature” that uniquely identifies each individual in the human species. Recently, the quest for identifying reliable markers of human identity has expanded into the field of neuroscience. A seminal work^1^ in this research area has highlighted that the expression of an individual’s brain connectome^2^ can act as a “fingerprint” that uniquely identifies a given individual among a large population of individuals solely on the basis of its brain connectome profile. This work^1^, along with others,^3,4^, laid the foundation for a new field that has taken the name of “brain fingerprinting” and, since then, its scope has rapidly expanded thanks to the fact that brain fingerprints can now be derived from structural magnetic resonance imaging (MRI)^5–7^, functional MRI (fMRI)^1,3,4,8^, electroencephalogram (EEG)^9–11^, or functional near-infrared spectroscopy (fNIRS)^12^, and they can also be related to behavioral and demographic scores ^13–17^. Methodologically, most of these works are based on extracting fingerprints from inter-individual functional connectivity profiles, also known as functional connectomes (FCs), that are understood as being the statistical dependence between spatially distinct regions^18^.

Only very recently has the fingerprinting field started to capitalize on the spatiotemporal complexity of fast neurophysiological signals recorded from magnetoencephalography (MEG) in order to investigate neural features of individual differentiation^16,19–22^. There are several reasons for doing so, since recorded MEG signals contain extremely rich information^23,24^. First, MEG signals measure direct cortical activity with a high temporal resolution as opposed to fMRI that only provides information about slow hemodynamic fluctuations. Second, the measured signals oscillate at multiple frequencies that allow for band-specific interpretations; and third, oscillations that resonate at different frequencies have a biological meaning that is related to cognitive functioning. Indeed, recent studies taking advantage of spontaneous electrophysiological recordings have provided new insights into the neurophysiological nature of brain fingerprints in healthy^16,19^ and clinical populations^20–22^.

So far, these studies have only focused on characterizing individual MEG signatures from task-free conditions, under which individuals are not engaged in any particular task^25,26^. However, resting-state activity does not capture the full range of interindividual differences in the functional organization of the brain^27,28^, nor can it fully predict brain-behavior relationships ^17,28–31^. Specifically, spontaneous brain activity fails to capture the functional reconfiguration of the brain that takes place as individuals engage in various activities^32,33^. Task-paradigms reliably perturb the ongoing dynamics of the core functional organization of the human brain, by modulating its connectivity patterns according to task demands and individualized responses^32–39^.

Hence, the next step is to explore whether individual signatures of identifiability from fast neurophysiological brain dynamics are modulated due to the task-dependent properties of the functional connectome. Since this is uncharted territory, there are several interesting aspects to be explored. How is individual identifiability affected by task-induced modulations? Are certain brain rhythms more specific for differentiation? How does the spatial organization of brain fingerprints —in terms of brain regions and systems—change with varying brain states? Finding an answer to these questions will enhance our knowledge of what the driving factors behind MEG connectome identification are. In this work, we addressed several of these questions by deriving brain connectivity fingerprints of MEG data from a cohort of individuals collected during several brain states. We started by estimating the functional connectomes of each individual in resting-state conditions, and three task-induced conditions. We found compelling evidence from whole-brain functional connectivity patterns that individuals were identified better when engaged in task conditions that were under the strict control of the experimenter (i.e, well-constrained), indicating that MEG connectome identifiability changes as a function of the task and its level of constraint. Notably, the contributions of brain regions and functional systems to individual identifiability were modified when engaged in task-induced brain states. In summary, the findings in this work indicate that the connectome fingerprint is not static, but is something that fluctuates and becomes more prominent while engaged in certain task-driven states. More importantly, we can track this fluctuating feature of connectome identifiability across several frequency components using direct neurophysiological signals captured by MEG. We hope that the findings reported in this work will provide new insights into the link between individual brain signatures and behavior.

## Results

We aimed to formally investigate three aspects of MEG signatures: *i)* How identification of individuals based on connectome features is affected by the brain state (as manipulated by environmental conditions); *ii)* To what extent the connectivity patterns needed for brain identification change as a function of the experimental design; *iii)* How task differentiability relates to individual brain fingerprints. A general scheme to investigate these aspects is illustrated in Figure 1. We explored MEG signatures of twenty individuals across a set of four different experimental conditions: resting-state (REST), narrative listening (PROSE), mismatch negativity (MMN), and Auditory Steady State Responses (ASSR) (Fig. 1a). The fingerprinting approach was applied to these MEG recordings, and started with estimating functional connectomes for each individual from test/retest MEG segments after source reconstruction (cf. Fig. 1a-b and see Methods for details). Next, the degree of differentiability for each condition was estimated using differential identifiability (Idiff) and success-rate (SR) metrics, computed from a mathematical object called identifiability matrix^4^ (Fig. 1c; see Methods for details). This identifiability matrix encodes the similarity of each individual with themselves (*Iself*; diagonal elements) as opposed to others (*Iothers*; off-diagonal elements), and *Idiff* conceptualizes the extent to which individuals were more similar to themselves than others^4^ (i.e., the difference between the average *Iself* and *Iothers* values). In addition, SR^1^ was used as a complementary score that provides the proportion of correctly identified individuals. Finally, we explored the *spatial specificity* of MEG fingerprints by estimating the degree of distinctiveness of each FC-edge for individual and task differentiability using intraclass correlation (ICC; Fig. 1d and see Methods for details). Given that MEG recordings are rich multi-spectral signals, the fingerprinting analysis was repeated for five typical frequency bands, as a means to identify which brain rhythms were most specific for individual differentiability across varying tasks.

**Figure 1.**
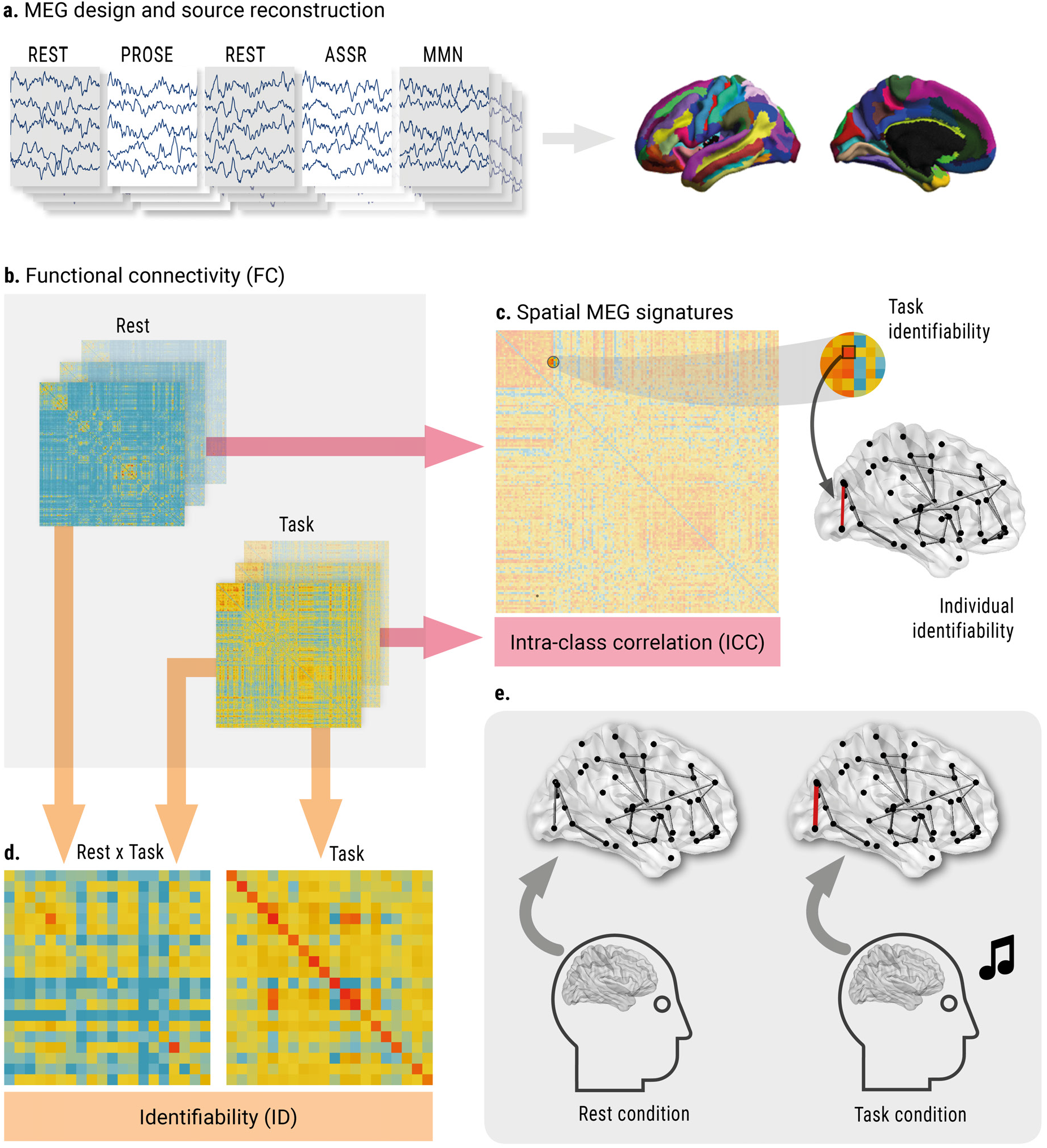
Exploring MEG signatures in naturalistic and neurophysiological states. **(a)** MEG signatures were explored in a set of recorded naturalistic and neurophysiological brain states *(left)*: Resting-state (REST; two sessions), Narrative Listening (PROSE), Auditory Steady State Responses (ASSR) and Mismatch Negativity (MMN). The recordings for each individual were preprocessed and source reconstructed to obtain a cleaned time series from each region of the Destrieux Atlas^40^ *(right)*. **(b)** Individual FCs from test/retest segments were obtained by using the functional connectivity measure of Amplitude Envelope Correlation (AEC) between all pairwise orthogonalized time series of the 148 regions of the Destrieux Atlas^40^. **(c)** The degree of differentiability in each environmental condition was derived from a mathematical object called identifiability matrix, which summarizes the degree of similarity between test FCs vs. retest FCs. **(d)** The *spatial specificity* of MEG signatures was assessed using edgewise intraclass correlation (ICC)^41^. This method was used to estimate the distinctiveness of each FC-edge for differentiating between individuals (individual identifiability), and differentiating between the set of environmental conditions (task identifiability). **(e)** The workflow from (a-d) allowed us to explore several aspects of MEG signatures derived from functional connectomes. First, to what extent identifiability changes as a function of task-induced brain states, by evaluating the degree of differentiability computed in (c). Second, to what extent the spatial specificity of fingerprints at the individual level changes across brain states as measured by the edgewise-ICC metric in (d). And third, whether we can identify a spatial signature that differentiates between the set of brain states (task identifiability) using edgewise ICC (d).

### Individual identification from MEG functional connectomes shows task-dependent aspects

We found that identifying individuals based on their functional MEG connectomes was better when they were engaged in explicit tasks (Fig. 2). Figure 2a reports the exemplary identifiability matrices for the alpha and beta frequency bands, whereas the matrices for the other frequency bands are reported in Supplementary Fig. 1. On average across all frequency bands, differentiation scores were lower for REST (SR 60.0%, Idiff 17.8%) compared to PROSE (SR: 74.5%, Idiff: 26.2%), ASSR (SR: 99.5%, Idiff: 37.2%) and MMN (SR: 100%; Idiff: 36.2%). This observation was confirmed using Non-parametric Wilcoxon rank tests between the individual Idiff scores of the rest and task states that were all statistically significant (*P*-values < 0.001, after Bonferroni correction for multiple comparisons). Notably, there was a drop in identifiability for combinations between “task-rest” states, where performances across frequency bands varied between 25.0%-63.0% for SR, and between 8.3%-18.5% for Idiff. These findings together indicate that individuals tend to be more differentiable during tasks than during rest. The results were robust to evoked fields’ effects as differentiation scores were computed from residualized task FCs (i.e., after regressing out the average signal across trials), and differentiability scores were not substantially altered when running the same procedure using non-residualized task FCs (Supplementary Fig. 2).

**Figure 2.**
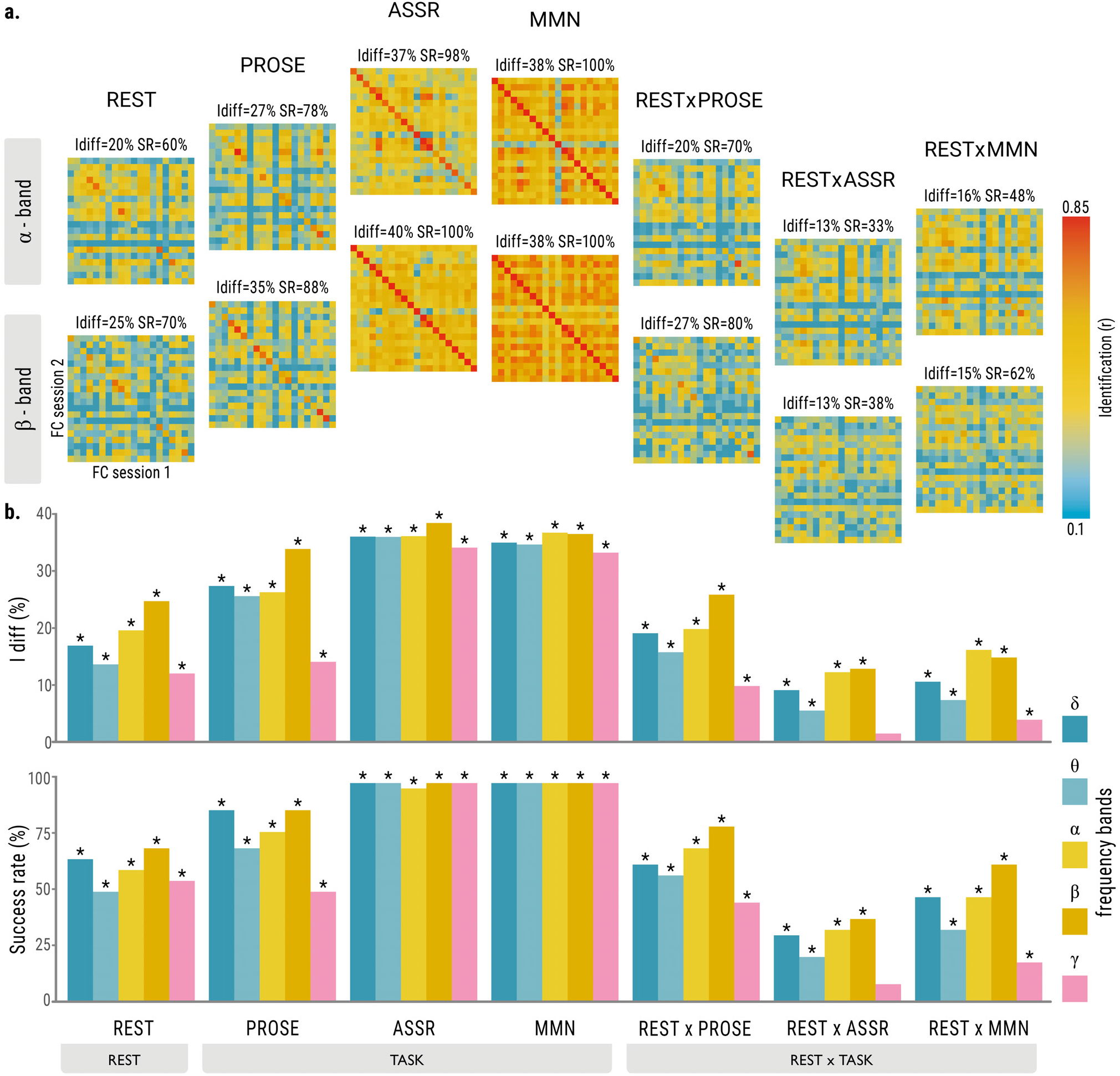
Task matters: identifiability scores across brain states. The figure illustrates the differential identifiability (Idiff) and success-rate (SR) scores across the brain states of resting-state (REST), task-based states (PROSE, ASSR, MMN), and combinations of rest and task brain-states (REST × PROSE, REST × ASSR, REST × MMN). **(a)** Identifiability matrices for each of the brain states for the alpha (8-13 Hz) and beta frequency bands (13-30 Hz) (results on the other bands are reported in Supplementary Fig. 1). **(b)**Bar plots summarizing the identification scores across the different brain states and frequency bands. The asterisks on top of the bar plots denote a significant identification score after permutation testing (see Methods for details on the null model employed).

An interesting observation is that in the PROSE state the delta and beta rhythms were most specific for individual connectome identification, whereas in the MMN/ASSR states no particular frequency range was specific, i.e., identification scores were consistent across frequency bands (Fig. 2 and Supplementary Fig. 1). Notably, the differentiation scores for the cross-state setting of REST x PROSE were similar to those of REST, which was not the case for the cross-state settings of REST × MMN and REST × ASSR (Fig. 2b). These observations might in large part be related to the degree of constrainedness of the task. In traditional task paradigms (MMN/ASSR) that are considered as overly constrained states, overall variability in connectivity is constrained (i.e., reduced noise). Conversely, REST is a totally unconstrained state and narrative listening (PROSE) is a more naturalistic paradigm that is less constrained than traditional task paradigms. In addition, the *Iother* and *Iself* elements of the identifiability matrices further suggest that the nature of brain state influences identifiability at the individual level, as their values increase and their distributions become narrower as a function of how constrained the brain state is (Fig. 3). In other words, the more constrained an experimental condition is, the more the within- and between-individual variabilities are reduced across test-retest sessions, which leads to increased fingerprinting levels by preserving or enhancing important individual signatures of connectivity.

**Figure 3.**
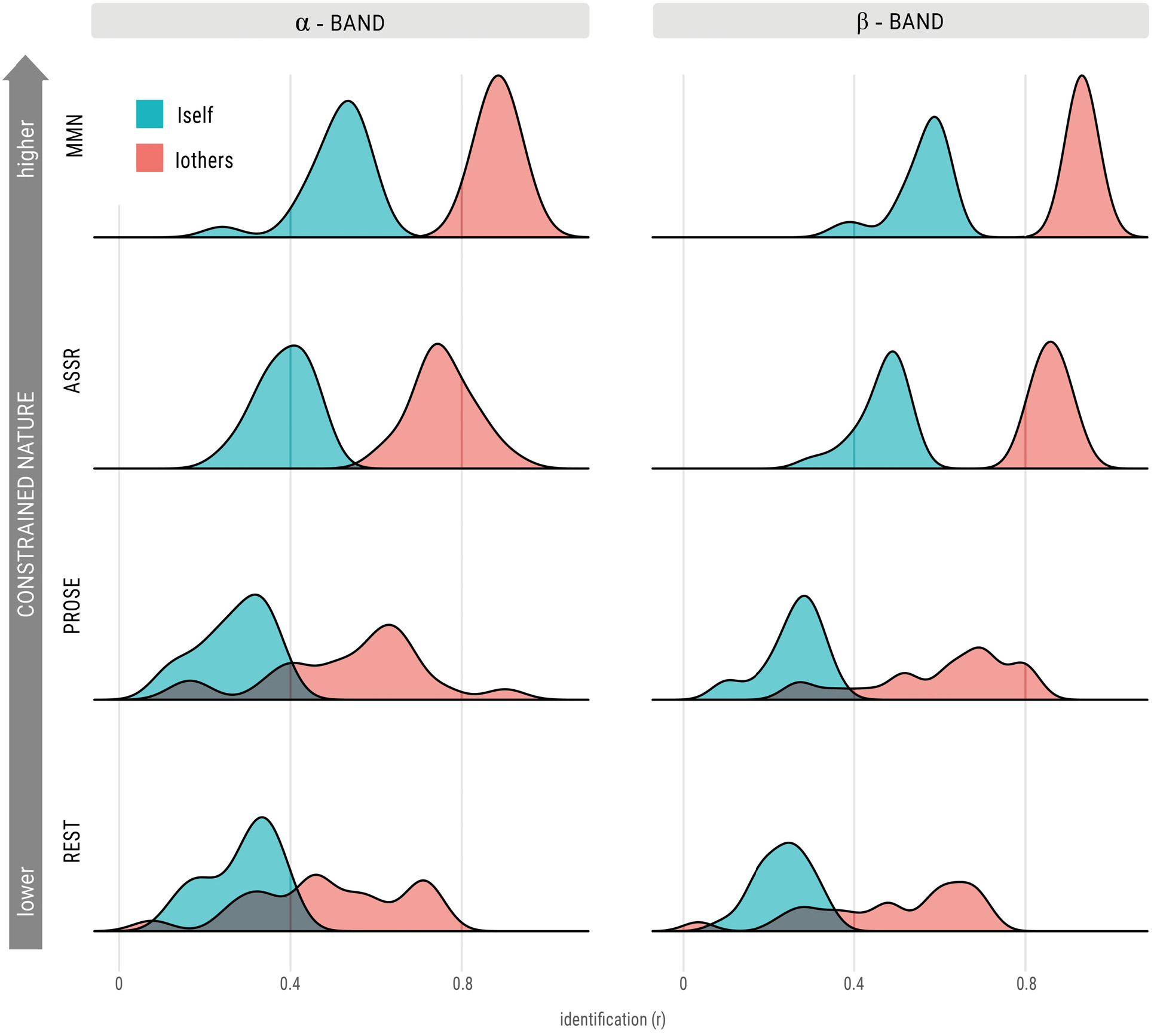
Connectome identification is dependent on the constrained nature of the brain state. Distributions of the *Iself* and *Iothers* values in the alpha and beta frequency bands for all brain states. The distributions indicate that within connectome similarity (*Iself*), and between connectome similarity (*Iothers*) change as a function of the constrained nature of the brain state. The *Iself/Iothers* distributions shift rightward and become narrower from unconstrained (REST) to well-constrained states (MMN/ASSR).

### Spatial MEG signatures of individual differentiation change according to brain state

Given that the identification rates computed at the whole-brain level do not provide information on the functional edges that contribute most towards individual identification, we used the edgewise Intraclass correlation metric (ICC; see Methods) to assess the spatial specificity of individual MEG signatures. We found that the spatial signatures of the most identifiable edges for individual differentiation were modified across brain states (Fig. 4-5a for alpha (8-13 Hz) and beta frequencies (13-30 Hz), other frequency bands reported in Supplementary Fig. 3-5). Specifically, for all frequency ranges we observed changes in the amount of reliable FC-edges for task-based states compared to the resting-state (Fig. 4-5a and Supplementary Fig. 3-5a). Similar results were obtained when refining the spatial exploration and looking at the regional counterpart of the ICC profiles across well-defined functional systems^42^ (Fig. 4-5b and Supplementary Fig. 3-5b), or at their nodal strength (i.e., taking the column-wise mean of the ICC matrices; Fig. 4-5c and Supplementary Fig. 3-5c).

**Figure 4.**
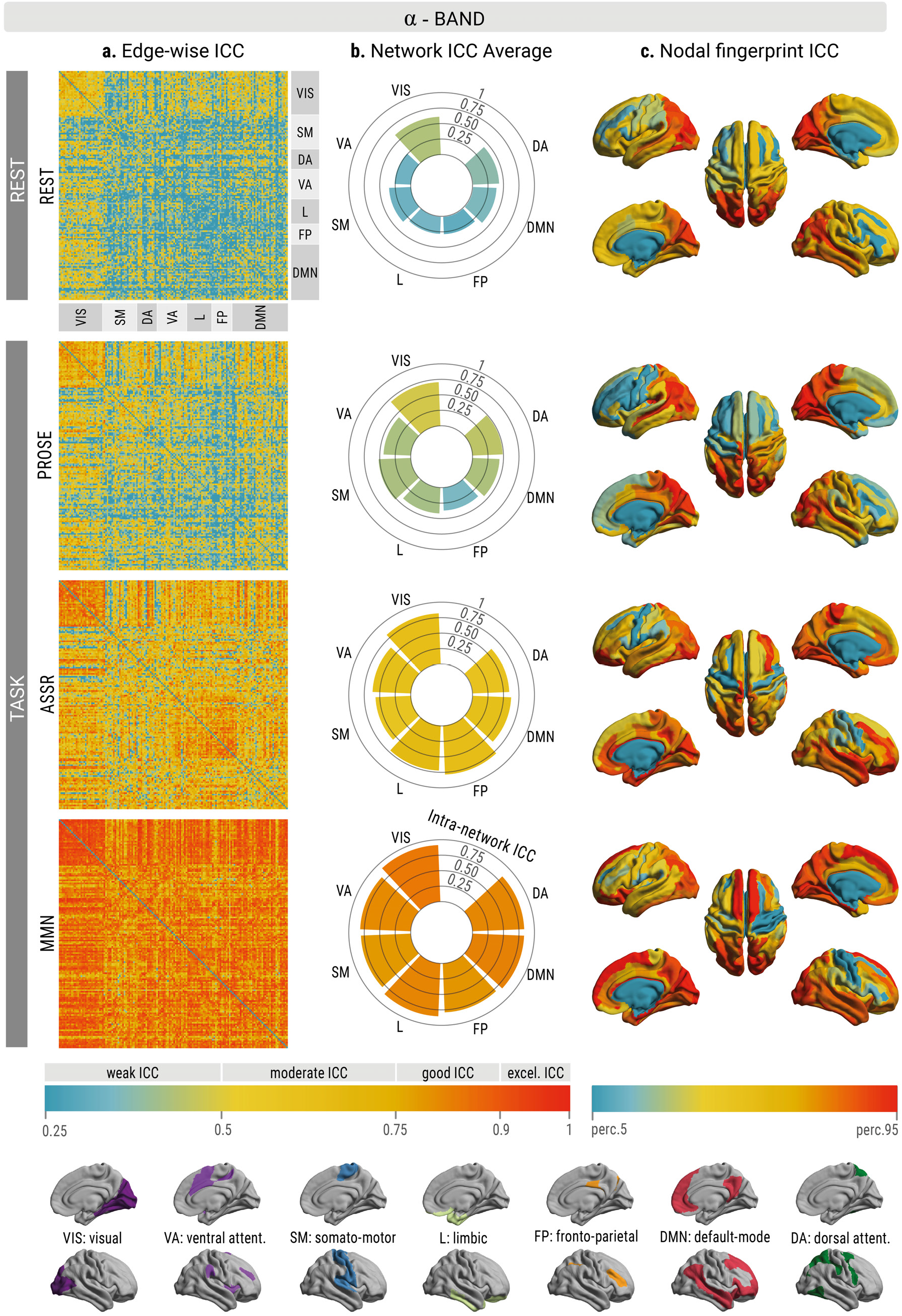
Spatial signatures of individual differentiability in the alpha frequency band. **(a)** Edgewise individual differentiability as measured by intraclass correlation (ICC) for each brain state for the alpha frequency band. The ICC values for each of the functional connections per brain state are shown. The higher the value, the more the connection is able to separate an individual from others in the cohort. The brain regions are ordered according to the seven intrinsic functional system organization proposed by Yeo and colleagues^42^. **(b)** The edgewise ICC scores are averaged within (axis) and between (color) all seven functional systems to better visualize fingerprint patterns within and between functional systems across brain states. **(c)** Nodal representations of the brain regions involved in individual differentiability during a specific brain state, represented at the 5-95th percentile threshold. The nodal strength of the ICC matrix (i.e., taking the column-wise mean) was used to characterize how central each brain region is for individual differentiation. *Abbreviations of Yeo’s functional systems* VIS = visual; SM = sensorimotor; DA=dorsal attention; VA=ventral attention; L= limbic; FP= frontoparietal; DMN= default mode network.

**Figure 5.**
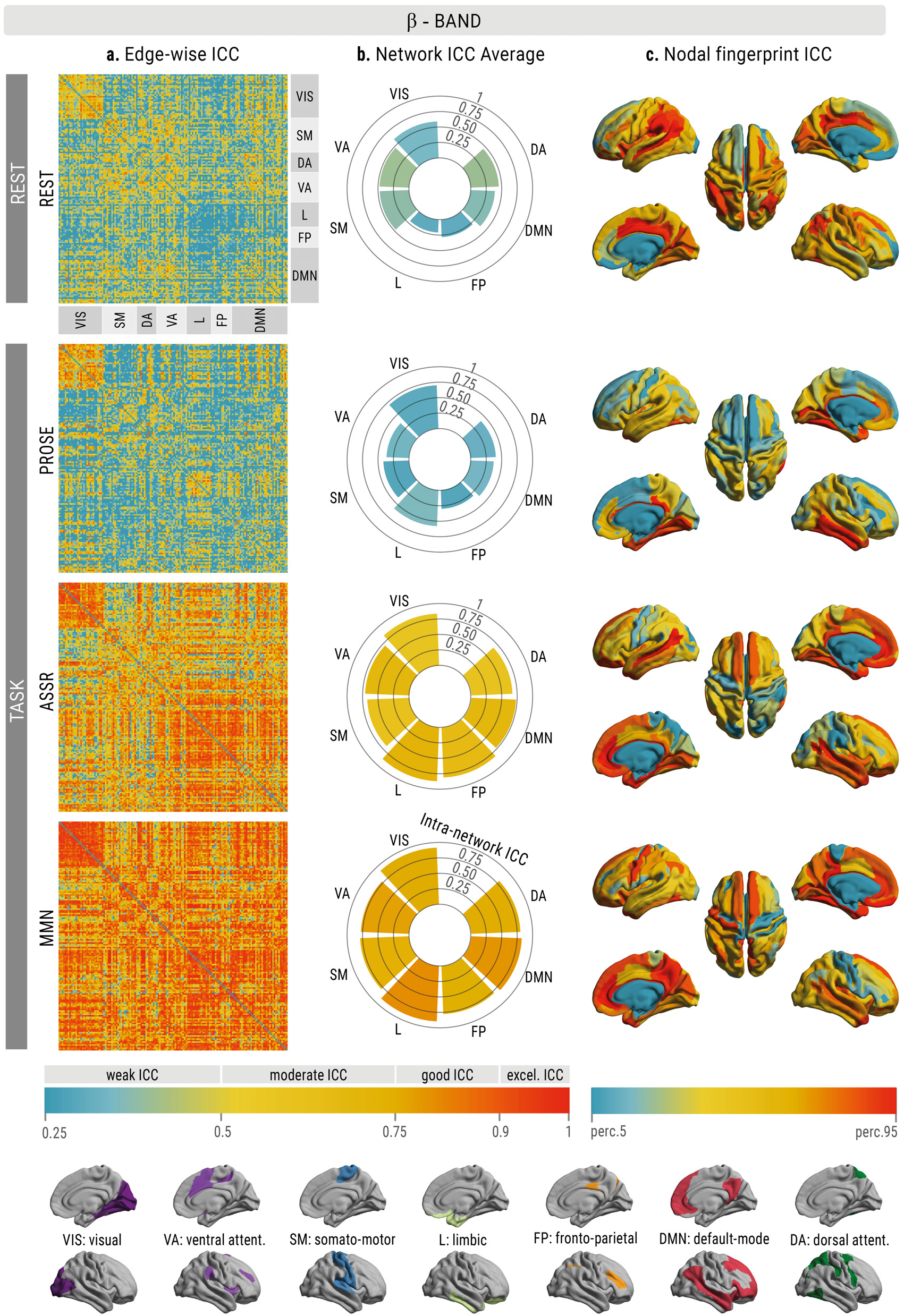
Spatial signatures of individual differentiability in the beta frequency band. **(a)** Edgewise individual differentiability as measured by intraclass correlation (ICC) for each brain state for the beta frequency band. The ICC values for each of the functional connections per brain state are shown. The higher the value, the more the connection is able to separate an individual from others in the cohort. The brain regions are ordered according to the seven intrinsic functional system organization proposed by Yeo and colleagues^42^. **(b)** The edgewise ICC scores are averaged within (axis) and between (color) all seven functional systems to better visualize fingerprint patterns within and between functional systems across brain states. **(c)** Nodal representations of the brain regions involved in individual differentiability during a specific brain state, represented at the 5-95th percentile threshold. The nodal strength of the ICC matrix (i.e., taking the column-wise mean) was used to characterize how central each brain region is for individual differentiation. *Abbreviations of Yeo’s functional systems* VIS = visual; SM = sensorimotor; DA=dorsal attention; VA=ventral attention; L= limbic; FP= frontoparietal; DMN= default mode network.

In addition, and in line with our previous analysis (Fig. 2), we found that the spatial signatures of individual differentiation varied as a function of the constrained nature of the task (Fig. 4-5; Supplemental Fig. 3-5). For the well-constrained states (MMN/ASSR), we observed a rigid spatial profile for all frequency ranges, whereas for the un/less constrained REST/PROSE states, we observed variability in the spatial profiles across frequency bands. Specifically, for alpha connectivity in REST the visual functional subsystem was the most specific hub for individual identification, whereas for PROSE the visual and somatomor subsystems were the most distinctive among individuals. For the beta-band connectivity, posterior regions belonging to the default mode and ventral attention systems were hubs of individual differentiation in the REST state, whereas in the PROSE state discrimination was mainly driven by visual and limbic regions. In contrast, for the MMN/ASSR states, a spatial profile consisting of regions belonging to higher-order systems (i.e., default mode, ventral attention, limbic) that span both posterior and frontal brain regions was most specific for all frequency ranges. Taken together, these findings show that spatial profiles of individual MEG signatures contain task-dependent aspects and that the nature of the task influences the spatial pattern across oscillatory rhythms.

### A spatial MEG signature differentiating brain states

After identifying the spatial MEG signatures that distinguish individuals, we explored whether there was a spatial pattern of connectivity edges that was able to differentiate among the set of brain states (i.e., task differentiability). The analysis of task differentiability was inspired by a previous fingerprint study^4^ and is also based on the edgewise ICC (see Methods). However, the ICC values should be interpreted in a different manner compared to individual differentiability. In this case, the higher the ICC value of an edge, the more distinctive it is for differentiating between the set of brain states across individuals. Results showed a specific spatial signature of functional subsystems that differentiates among brain states (Fig. 6). In particular, the edges within the limbic, somatomotor and default mode functional systems were most involved in the spatial signature of task differentiability, whereas inter-system connectivity was less involved. At the nodal level, regions with the highest ICC strengths mainly spanned the temporal and frontal lobes, including the somatomotor regions.

**Figure 6.**
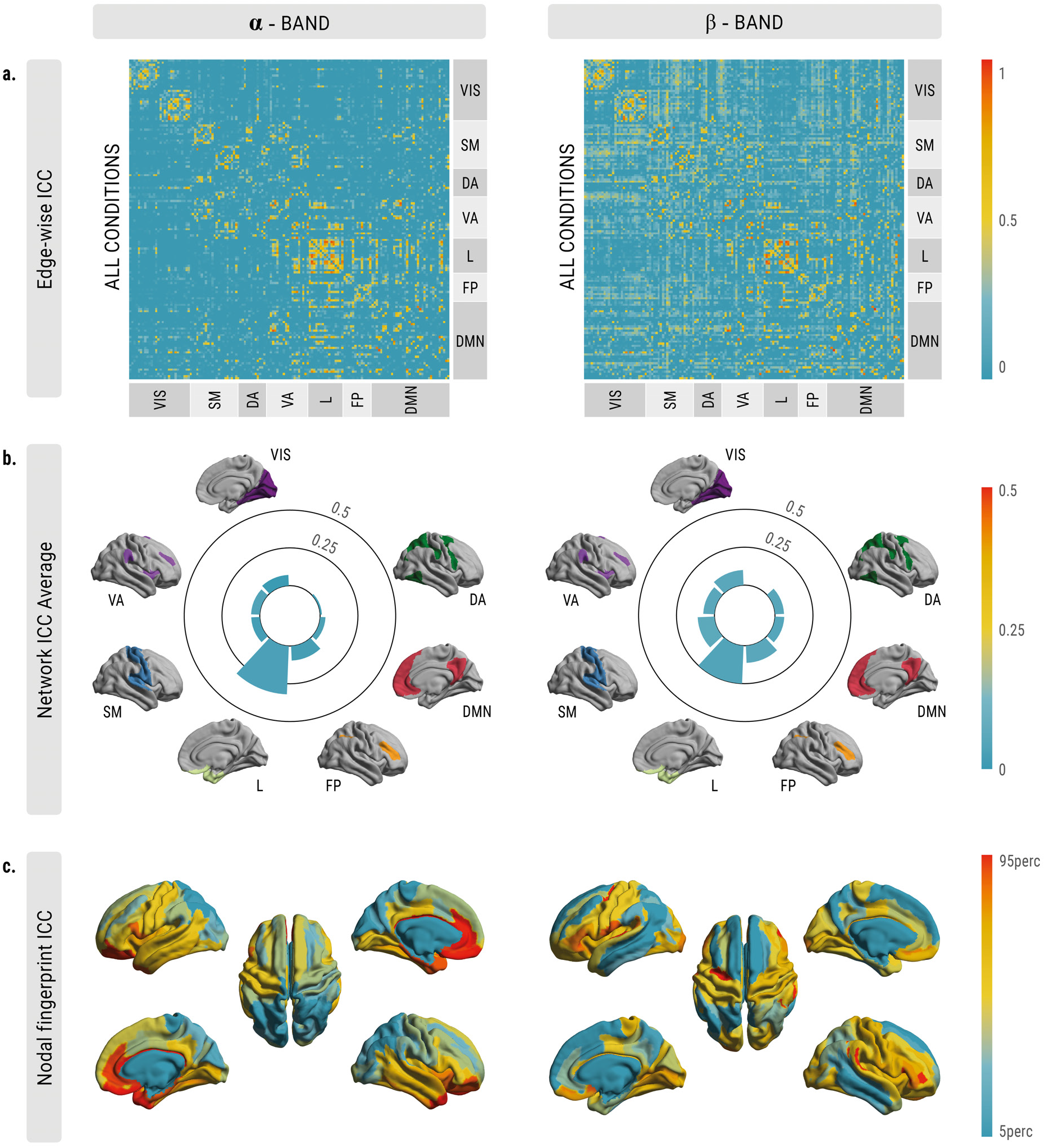
Spatial signature of task differentiability. **(a)** Edgewise task differentiability as measured by intraclass correlation (ICC) for the alpha and beta frequency bands. The higher the ICC value, the more the connection is able to differentiate between the set of brain states. The brain regions are ordered according to the seven intrinsic functional system organization proposed by Yeo and colleagues^42^. **(b)** The edgewise ICC scores are averaged within (axis) and between (color) functional systems to better visualize fingerprint patterns. **(c)** Nodal representations of the brain regions involved in differentiating between brain states, represented at the 5-95th percentile threshold. The nodal strength of the ICC matrix (i.e., taking the column-wise mean) was used to characterize how central each brain region is for task differentiation. The analysis shows that the connections of the limbic, somatomotor and anterior default mode functional systems contributed most to the fingerprint profile of task differentiability.

## Discussion

Connectivity patterns from neural signals captured with MEG can differentiate individuals within a cohort, similarly to fingerprints. Brain fingerprints characterized from resting-state neurophysiological activity are predictive of phenotypic measures (e.g., age)^16,19^, and improve our understanding of brain dysfunction^20,22^. While individuals can be reliably identified from resting-state MEG dynamics, it is not known to what extent we can differentiate individuals from a range of fast brain signals when they are explicitly engaged in task-related behavior. This relates to the key question: does the identifiability of the MEG functional connectome change as a function of tasks?

Here, we explored several aspects of MEG signatures in three directions i.e., to ascertain: *i)* Whether the individual identifiability of functional connectomes changes with task-induced manipulations, *ii)* whether spatial MEG signatures demonstrate spatial variability along with a change in brain state, and *iii)* whether task-induced brain states can be identified from functional connectomes. Using the connectome fingerprinting procedure, we found that identifiability scores improved when individuals were engaged in task-states relative to the resting-state (Fig. 2), providing compelling evidence for a task-dependent axis of MEG fingerprints. These findings suggest that even though task-induced changes in functional connectivity are small perturbations of a stable intrinsic network architecture^32,34,35^, they enhance individual differences in neural circuitry^43,44^, which in turn increases the identifiability of individuals’ connectomes. Indeed, the presence of an intrinsic spatial organization of ongoing oscillatory signals has been recently reported^45^, and accordingly our observations show for the first time that state-dependent modulations of this intrinsic organization are functionally relevant for individual connectome identification.

Using a design with data acquired both during task execution and rest ensures that the improvements in individual differentiability from task-relative to resting-state derives from task-induced changes in FCs and individual differences in these modulations. In addition, the fingerprinting scores for the MMN/ASSR states cannot be explained by basic features of the tasks such as the sensory modality, rate and timing of stimulus presentation, since we regressed out task-activity, and the individual differentiability was similar when performed with a more conservative connectome fingerprint analysis on the raw time courses (Supplementary Fig. 2). This suggests that those basic features that differ among the neurophysiological task paradigms did not hinder, nor improve the fingerprinting performances, and therefore were not a contributing factor to the high identification scores.

What other factors influence individual identifiability? According to the findings reported, individual identifiability is also determined by the constrained nature of the task (Fig. 2-3, Supplementary Fig. 1). We found that in well-constrained tasks identifiability scores are high and coherent across frequency bands. This can be attributed to the fact that well-constrained tasks offer a strict controlled manipulation of the brain state that taps into relevant neural circuitry^46^, and amplifies any individual differences occurring above the common task-circuity^43,44^. In contrast, in less-constrained tasks identifiability is lower and shows specificity for certain frequency ranges. On the one hand, in the case of the resting-state this can largely be explained due to its totally unconstrained nature^25^, whose connectivity patterns in this state are associated with reduced within-individual test-retest stability over multiple recordings, due to the influences of a mixture of processes that are not easy to quantify, such as arousal, attention and mind wandering^44^. On the other hand, the narrative listening condition is a compromise between unconstrained and well-constrained states, since it introduces some boundaries to mental activity through an ecological valid stimulus (audio fragment) that is similar to real life situations^47^. Yet, the PROSE state is influenced by a mixture of processes that reduces the stability of connectome similarity both within and across individuals. In other words, the choice of the task paradigm is important for identifying individuals from their functional connectomes. This might have implications for precision medicine, as choosing the appropriate environmental setting could improve the link between features of connectivity and individual phenotype scores (e.g., clinical outcomes, fluid intelligence, etc.).

The individual specificity of FCs in the delta and beta frequency ranges during narrative listening is conceptually in agreement with previous investigations of MEG activity recorded during this task. Both delta and beta activities emerge in the literature as subserving processes in language comprehension, and one could therefore speculate that these MEG signatures capture representations of language comprehension^48^ that are specific to each individual. Conversely, for well-constrained tasks there is no direct relationship of identifiability being salient in a particular frequency range, in contrast with previous studies which report modulations in theta^49,50^ (MMN), and gamma^51^ (ASSR) frequencies.

Similar to individual identifiability computed at the whole-brain level, individual spatial MEG signatures are modified according to the constrained nature of the task (Fig. 4-5 and Supplementary Fig. 3-5). For well-constrained tasks, the higher-order and limbic functional subsystems acted as a core signature for identification across all frequency bands. In the less-constrained tasks of rest and narrative listening, the spatial signatures of individual differentiation varied across frequency ranges. For instance, in the alpha and beta bands, the functional connections within the visual and limbic systems contributed the most to identifiability (Fig 4-5), while connections of other subsystems were most prominent for the delta, theta and gamma frequency bands (Supplementary Fig. 3-5). These observations show some correspondence to fMRI results, in which the main drivers of functional connectome identifiability reside in areas related to higher-order cognitive functions, such as the frontoparietal and default mode functional subsystems^1,4^. Notably, we find some spatial divergence in the fingerprint patterns relative to fMRI literature. This is not surprising since the nature of brain signals measured by both modalities is quite different, and the relationship between hemodynamic and electrophysiological connectivity in response to task-demands is largely unknown^52^. Note that our work does not directly address the cross-modality difference of connectome fingerprints. Future works that compare identical brain states in both modalities will be better suited to find out whether fingerprinting patterns induced by distinct tasks are shared across neuroimaging modalities.

Based on previous work, we wondered if MEG specific connections could differentiate between brain states *regardless* of the individual. Our results indicate that this is the case, and that a spatial task-differentiability signature mainly consists of connections within the limbic, default mode, and somatomotor functional subsystems (Fig. 6). These subsystems are the ones in which we observed the greatest variation when transitioning across the different tasks employed, in line with the notion of the brain’s “functional reconfiguration” across tasks^37^. This technique could be used to select and extract the connectivity features that mostly differentiate between brain states, in health and disease. Future work should further explore and exploit this possibility.

This work comes with some considerations and limitations. First, on the basis of previous findings it is known that the choice of connectivity measures to derive the FCs is a factor that influences fingerprinting in MEG^19^. In addition, identifiability could be susceptible to the choice of brain atlas and the latter’s role should be further identified. However, other factors such as typical recording artifacts (head motion, heartbeats, eye movements, etc.), that might be representative of individuals do not seem to confound individual identification from MEG recordings^16^. Furthermore, signatures derived from MEG are robust to the information of participants’ anatomical head-position that is embedded in the MEG source imaging kernels, as it has been shown that this information is not sufficient to uniquely drive identification of individuals^16^. Comprehensive future studies will be needed to clarify the impact of several factors on MEG fingerprints during task-induced manipulations such as connectivity measures, choice of MEG source modeling, and parcellation schemes. We look forward to future work in electrophysiological fingerprinting that confirms and expands the present findings. Second, electrophysiological recordings from limbic regions are known to be affected by artifacts, and the accuracy of MEG in detecting signals in deeper cortical regions is still being debated^53^. As such, the findings reported on the involvement of the limbic subsystem in spatial signatures of task and individual differentiability should be interpreted with care.

Finally, the unique design of the employed dataset limited the sample size of the current work.

Therefore, the prediction of brain-behavior relationships from MEG fingerprints was not investigated, as recent studies have demonstrated that these predictions are not reliable with small sample sizes^54,55^. However, previous works have proven that fingerprints derived from functional connectomes of resting-state dynamics are predictive of individual differences in behavioral phenotypes^16,19^. Since our findings indicate task-dependent aspects of MEG signatures, we could speculate that individual differences that are amplified by tasks will improve the prediction of underlying related phenotypes that are not detectable by solely investigating the resting-state. Therefore, connectome fingerprinting from MEG signals during task conditions could help to improve the resolution and characterization of robust brain-behavior relationships.

To conclude, our work shows a task-dependent axis of brain fingerprints derived from fast electrophysiological signals, highlighting that task-induced brain states amplify meaningful interindividual differences in functional connectivity. In particular, individual identifiability increases when the brain state is driven by a well-constrained task compared to resting-state.

## Methods

### Participants and data acquisition

All data collection was performed at the MEG Laboratory of IRCCS San Camillo Hospital, Venice, Italy. Participants were recruited on a voluntary basis, upon signing a written consent. Participants had a mean age of 29.1 (SD = 5.82) years and on average 17.05 (SD = 2.50) years of education. Fifteen out of 20 participants were female. All participants reported no auditory issues. Before entering the magnetically shielded room, participants underwent initial preparation, which consisted of the placement of three head coils, to monitor head position during MEG recording, and six additional electrodes. These electrodes were used to record VEOG, HEOG, and ECG with bipolar montage. After the coils were positioned, coil positions and head shape were digitized using a Polhemus Isotrak system. Continuous MEG signals were acquired using a whole head 275-channel system (CTF-MEG). MEG data were collected with a sampling rate of 1200 Hz, with a hardware anti-aliasing low pass filter at 600 Hz. During the recordings participants remained in a seated position. The MEG session consisted of a series of recordings, all in a fixed order:

1. Eyes open resting-state - session 1 (REST; 5 min)
2. Narrative Listening (PROSE; 5 min)
3. Eyes open resting-state - session 2 (REST; 5 min)
4. Mismatch negativity (MMN; 3 min).
5. Auditory Steady State Responses at 40 Hz (ASSR; 6 min).

Details on each session are reported below.

### Resting-state eyes open

During the two sessions of resting-state participants were instructed to maintain visual fixation on a central crosshair, while avoiding excessive eye movements.

### Narrative Listening

During the narrative listening session, participants were asked to listen to 5 min audio recordings (a fragment from an audiobook “20’000 leagues under the sea”, by Jules Verne, read by a professional actor). All participants listened to the same fragment. To ensure people paid attention to the audio, they were initially informed that afterwards they would be asked some questions on the content of the recordings. These yes/no questions pertained to the content of the audiobook (e.g., was the story settled in the south pole), and were designed to check whether participants were paying attention during the recording.

### Mismatch Negativity

In this task participants were exposed to a series of tones, consisting of standard tones interleaved with deviant ones. Standard and deviant tones were generated as two tones of 500 Hz and 550 Hz (with 6 harmonics having 2,4,6,8,10, and 12 times the original frequency) lasting 200 ms. The sounds that acted as deviant and standard tones were counterbalanced across participants. To avoid that the deviant tones could be too close to each other during the task, we opted for a pseudorandom presentation of stimuli. The pseudorandom sequence was generated as follows. We first created a series of blocks of stimuli composed of several sounds: blocks consisting of three to eight standard stimuli, and blocks consisting of three to eight standard tones followed by a deviant one. In total, all blocks consisted of 300 stimuli, with 240 standard and 60 deviant sounds (hence, 25% of the stimuli were deviant). Afterwards, for each participant all blocks were shuffled to generate a unique sequence of sounds. Importantly, as deviant tones appeared only at the end of a block that started with three to eight standard tones, in the final (pseudo-random) sequence, deviant stimuli were always interspersed by at least three standard tones. Participants were not aware of the division in blocks and when performing the MMN task, sounds were presented as a stream of stimuli. In the original recording session two counterbalanced versions of the MMN were administered, one with an Inter Stimulus Interval (ISI) of 500 ms and one with an ISI of 3000 ms. For the aim of the present study, only the blocks with 3000 ms ISI were used, to ensure having epochs long enough to calculate the required connectivity matrices.

### ASSR

In this paradigm participants were exposed to a 1000 Hz tone, whose amplitude was modulated with a 40 Hz envelope. The same stimulus was used in previous publications by the research group^56–58^ and was proven to be able to elicit a 40 Hz entrainment, widespread in the brain, but mostly localized in the right auditory areas. The session consisted of 180 stimuli, each lasting 1 second, and with a fixed ISI of 1 second. Code used to generate the ASSR sounds can be found at: https://github.com/giorgioarcara/MEG-Lab-SC-code/tree/master/tDCS-ASSR.

#### MEG data preprocessing

MEG data preprocessing was performed using Brainstorm^59^ (version November 2018) in MATLAB 2016b (Mathworks, Inc., Massachusetts, USA), which is documented and freely available for download online under the GNU general public license (http://neuroimage.usc.edu/brainstorm). Continuous data were initially resampled at 600 Hz filtered with a notch (50 Hz and harmonics at 100, 150, 200 and 250 Hz) and a high pass filter at 0.1 Hz. Then the Signal-Space Projection algorithm (SSP) was used to identify and remove cardiac and eye movement artifacts from the recordings. For sessions with event-related responses (i.e., MMN and ASSR), triggers associated with stimulus presentation were used to segment continuous data into epochs. Digital triggers were adjusted off-line according to the actual acoustic stimulus presentation to improve accuracy of trigger timing.

For the source analysis, Individual T1 MRI scans were segmented by means of the recon-all routine of FreeSurfer^60^ image analysis suite, which is documented and freely available for download online (http://surfer.nmr.mgh.harvard.edu/). MRI and MEG data were registered according to the head-coil positions, identified with neuronavigation procedure. From the segmented MRI data, the MEG forward model was calculated with the Boundary Element Method (BEM). Source reconstruction was calculated on the cortex surface with the wMNE (weighted Minimum Norm) algorithm, using the Brainstorm default settings (with fixed source orientation, constraining the dipoles to be normal to cortex, using depth weighting with Order[0,1] = 0.5 and Maximal amount = 10; noise covariance regularization = 0.1, and specifying regularization parameter 1/λ by setting Signal-To-Noise Ratio = 3). The noise covariance was calculated from 3 minutes of empty room recording, made at the end of the recording session for each participant. Source time series were reconstructed into 148 cortical regions of interest (ROIs) according to the Destrieux atlas^40^ and dimension-reduced through the first principal component of all signals within each ROI using Principal Component Analysis (only the first component was retained). In addition, following a majority voting procedure each cortical region from the Destrieux atlas was assigned to one of the seven-resting state networks defined by Yeo and colleagues^42^. In order to make sure identifiability scores were not influenced by basic features of the task in MMN/ASSR conditions, task-activity in these conditions (i.e., mean across the epochs) was regressed out from the ROI time series in every epoch. Then, ROI source time series were divided into epochs of 8s duration and band-pass filtered into five commonly used frequency ranges (delta 1-4 Hz, Theta 4-8 Hz, alpha 8-13 Hz, beta 13-30 Hz and gamma 30-48 Hz). This length of epoch was chosen based on previously reported work that investigated the effect of epoch length on functional connectivity^61^.

#### Functional connectome generation

Functional connectomes were derived using the orthogonalized amplitude envelope correlation with spatial leakage correction (AEC)^62^. ROI time-series were first orthogonalized in the time domain with a pairwise leakage correction, before amplitude envelopes were determined by means of Hilbert transform to compute the corresponding Pearson correlations coefficients from all possible pairs, yielding 148 × 148 symmetric functional connectomes as a result. Two FCs (per frequency band, and individual) named test and retest FCs, were generated using test/retest MEG segments from each environmental condition. For the resting-state condition, the two separate acquired recordings were tagged as test and retest segments. For the task-conditions, the epochs in the first half of the session were tagged as test, and the epochs in the second half of the session as retest.

#### Fingerprinting and individual identifiability

Identifiability measures were obtained from the sets of test-retest FCs for each frequency band of interest. The identifiability metrics were computed within each condition (REST, PROSE, ASSR, MMN), and for 3 cross-conditions (REST × PROSE, REST × ASSR, REST × MMN). The methodology for the identifiability measures is inspired by recent work on maximization of connectivity fingerprints in human functional connectomes^4^. In this work, the authors proposed the ‘*differential identifiability*’ measure, which provides a robust continuous score of the fingerprinting level of a specific dataset. This measure is based on a mathematical object known as the ‘identifiability matrix’, which is a square and non-symmetric similarity matrix that encodes the information about the self-similarity of each individual with itself (*Iself*, main diagonal elements), and the similarity of each individual with the others (*Iothers*, off-diagonal elements) across the test-retest FCs. The similarity between the test-retest FCs was quantified as the Pearson’s correlation coefficient. The difference between the average *Iself* and *Iothers* values expressed in percentages is defined as the differential identifiability, and provides a robust group level-estimate of identifiability at the individual level from a specific dataset. The higher the *Idiff* score, the higher the individual differentiation in the cohort; the smaller the *Idiff* score, the more difficult it is to identify individuals from the cohort. Finally, we measured the Success-rate^1^ of the differentiation procedure as the percentage of individuals correctly identified out of the total number of individuals in the cohort. In other words, it expresses the percentage of cases with higher within- (*Iself*) vs. between-individuals (*Iothers*) FCs similarity. It is worth noting that in the present work average differentiation scores were reported for the cross-fingerprint setting of rest and tasks (across four possible combinations). Namely: 1) task-test FC vs. rest-test FC; 2) task-retest FC vs. rest-test FC; 3) task-test FC vs. rest-retest FC and; 4) task-retest FC vs. rest-retest FC.

In order to define the statistical significance of the obtained differential identifiability and success-rate scores, we performed a permutation testing analysis (1000 permutations)^19^. Specifically, for each iteration the identifiability matrices were randomly shuffled, before the measures of differential identifiability and success-rate were computed from the resulting surrogate identifiability matrix. A nonparametric ‘null’ distribution for success-rate and differential identifiability was then generated from all iterations. The *P*-values were computed as the proportion of times the permuted values of success-rate and differential identifiability exceeded those of the original scores.

#### Spatial specificity of individual and task MEG signatures

We derived the spatial specificity of the MEG signatures for each experimental condition and frequency band using edgewise intraclass correlation (ICC)^41^. Borrowing from previous work on identifiability^4^, we used ICC to quantify the edgewise reliability of individual connectomes. ICC is a widely used statistical measure that assesses the agreement between units (rating/scores) of different groups (raters/judges). The higher the ICC coefficient, the stronger the agreement between two observations^63^. Here, we used ICC to determine the *edgewise individual identifiability*, that quantifies the similarity between test and retest for each edge (i.e., functional connectivity value between two regions). In other words, the higher the ICC value on edge, the more consistent that edge’s value is within individuals between test and retest, and in turn, the higher the “individual fingerprint” of that edge. In addition, following the rationale of the ICC, we can also quantify the *edgewise task identifiability*, which quantifies the contribution of each edge towards separating the different environmental conditions across individuals. In this case, the tasks are considered as “raters”, and “scores” are given by individuals. Here, the higher the ICC, the more an edge can separate between the different tasks across individuals, and in turn, the higher the “task fingerprint” value of that edge. The resulting ICC matrices for both *the individual and task edgewise identifiability* were not thresholded and the ICC scores were interpreted according to the latest guidelines^64^; below 0.50: poor; between 0.50-0.75: moderate; between 0.75 and 0.90: good; and above 0.90: excellent. Finally, to explore the spatial organization of the individual-, and task MEG signatures we computed nodal fingerprinting scores (i.e., mean ICC over columns) from both the individual and task edgewise ICC matrices. This measure is an indication of the contribution of each brain region towards individual- or task identification.

## Data availability

Raw data are available from IRCCS San Camillo Hospital after formal requests and, if needed, after approval by the local Ethics committee for the intended use.

## Code availability

The code (in MATLAB) used for the analysis will be made available upon acceptance of the manuscript on EA EPFL webpage and a git repository. Code to generate the sounds of the ASSR task is available at: https://github.com/giorgioarcara/MEG-Lab-SC-code/tree/master/tDCS-ASSR.

## Acknowledgements

We would like to thank Diana Badder and Catherine Puntas for their help polishing the manuscript. This study was supported by a Ministry of Health Operating Grant to San Camillo Hospital IRCCS Venice (RRC-2021-23670183). GP was supported by GR-2019-12368960 from the Italian Ministry of Health. GA was supported by GR-2018-12366092 from the Italian Ministry of Health. E.A. acknowledges financial support from the SNSF Ambizione project “Fingerprinting the brain: Network science to extract features of cognition, behavior and dysfunction” (grant number PZ00P2_185716). We acknowledge the use of the Roboto and Roboto condensed open license fonts in the figures (Copyright 2022 Christian Robertson Licensed under the Apache License, Version 2.0).

