## Supplementary Materials for "Task matters: individual MEG signatures from naturalistic and neurophysiological brain states"

Supplemental Information

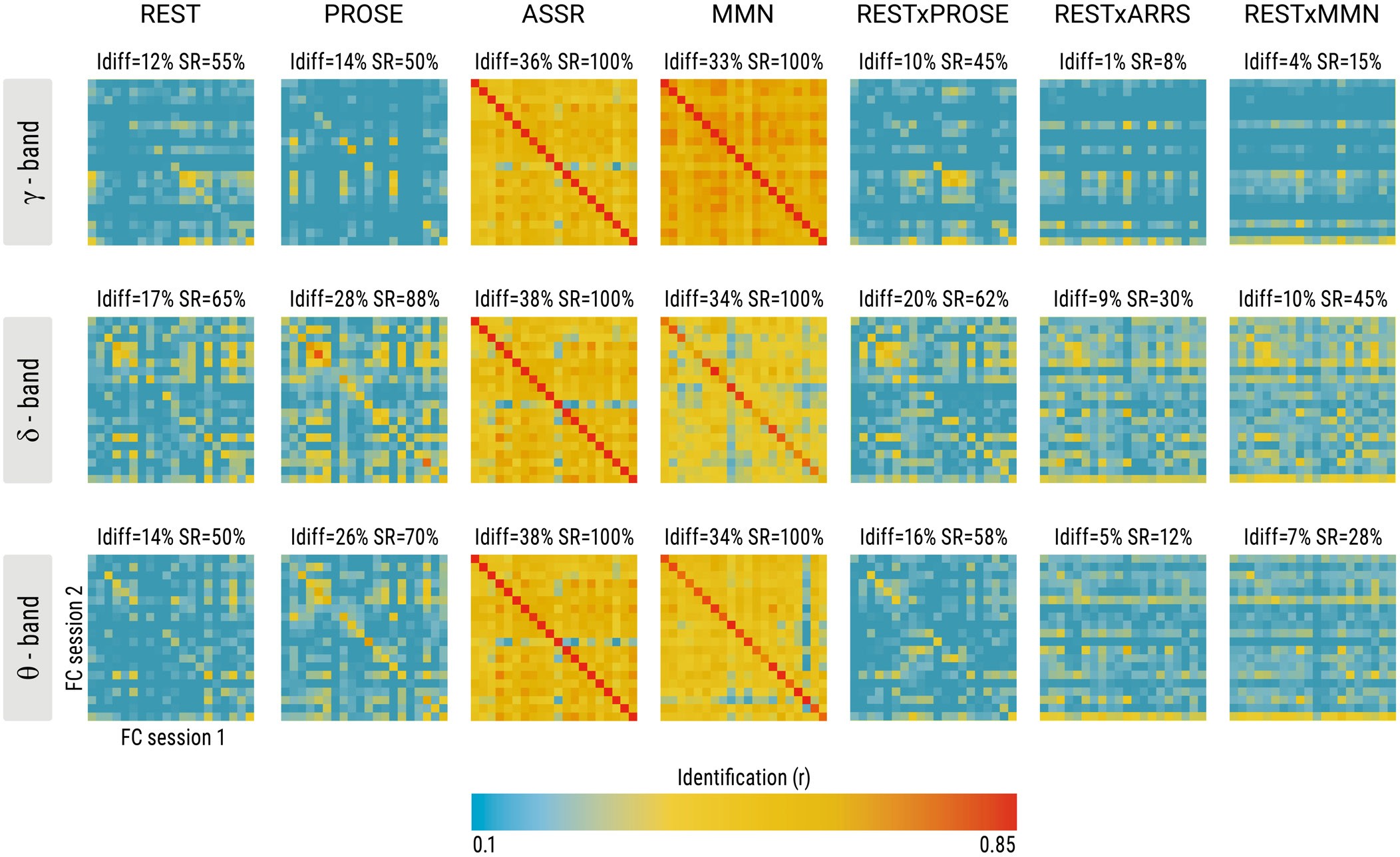

**Supplementary Figure 1 Identifiability scores of connectome fingerprinting across brain states in delta, theta and gamma frequency bands.** The figure illustrates the Identifiability matrices for each of the brain states for the delta, theta and gamma frequency bands. On top of each matrix two complementary scores of identifiability Idiff and SR are provided. Identifiability was computed for the following brain-states: resting-state (REST), task-based states (PROSE, ASSR, MMN) and the combination between rest and task brain-states (REST-PROSE, REST-ASSR, REST-MMN).

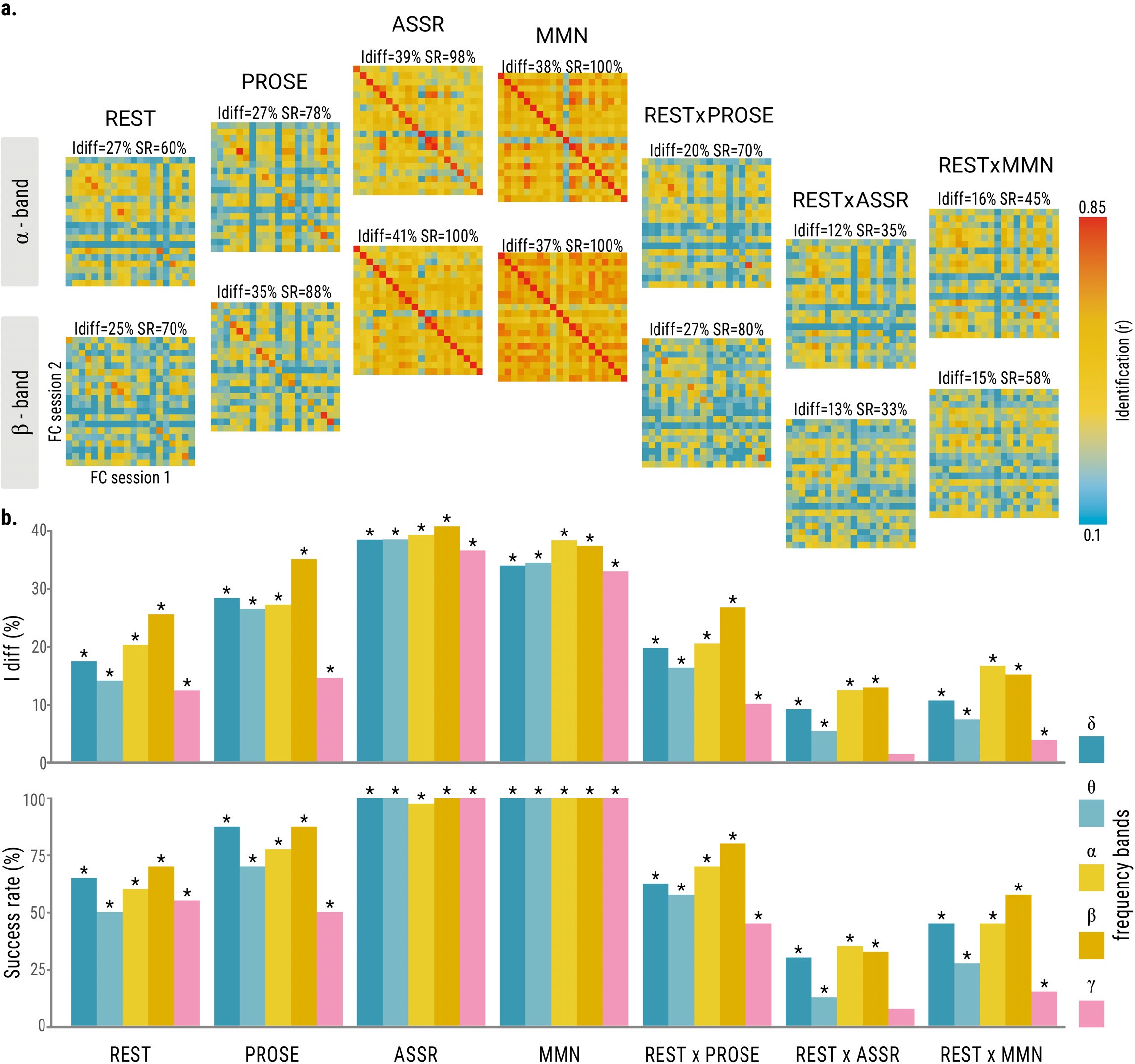

**Supplementary Figure 2 Identifiability scores of connectome fingerprinting based on raw-time courses for MMN/ASSR.** The fingerprinting procedure was applied on the raw-time series for the MMN/ASSR states, and were not corrected for mean task-activity (e.g. no task-regression). The figure illustrates the differential identifiability (Idiff) and success rate (SR); two complementary scores of the identifiability level across brain-states of resting-state (REST), task-based states (PROSE, ASSR, MMN) and combinations of rest and task brain-states (REST-PROSE, REST-ASSR, REST-MMN). **(a)** Identifiability matrices for each of the brain-states for the alpha and beta frequency bands. On top of each matrix the Idiff and SR scores are provided. **(b)** Bar plots showing the summary of the identification

metrics of Idiff and SR across the different brain-states and frequency bands. The asterisks on top of the bar plots denote a significant identification score after permutation testing.

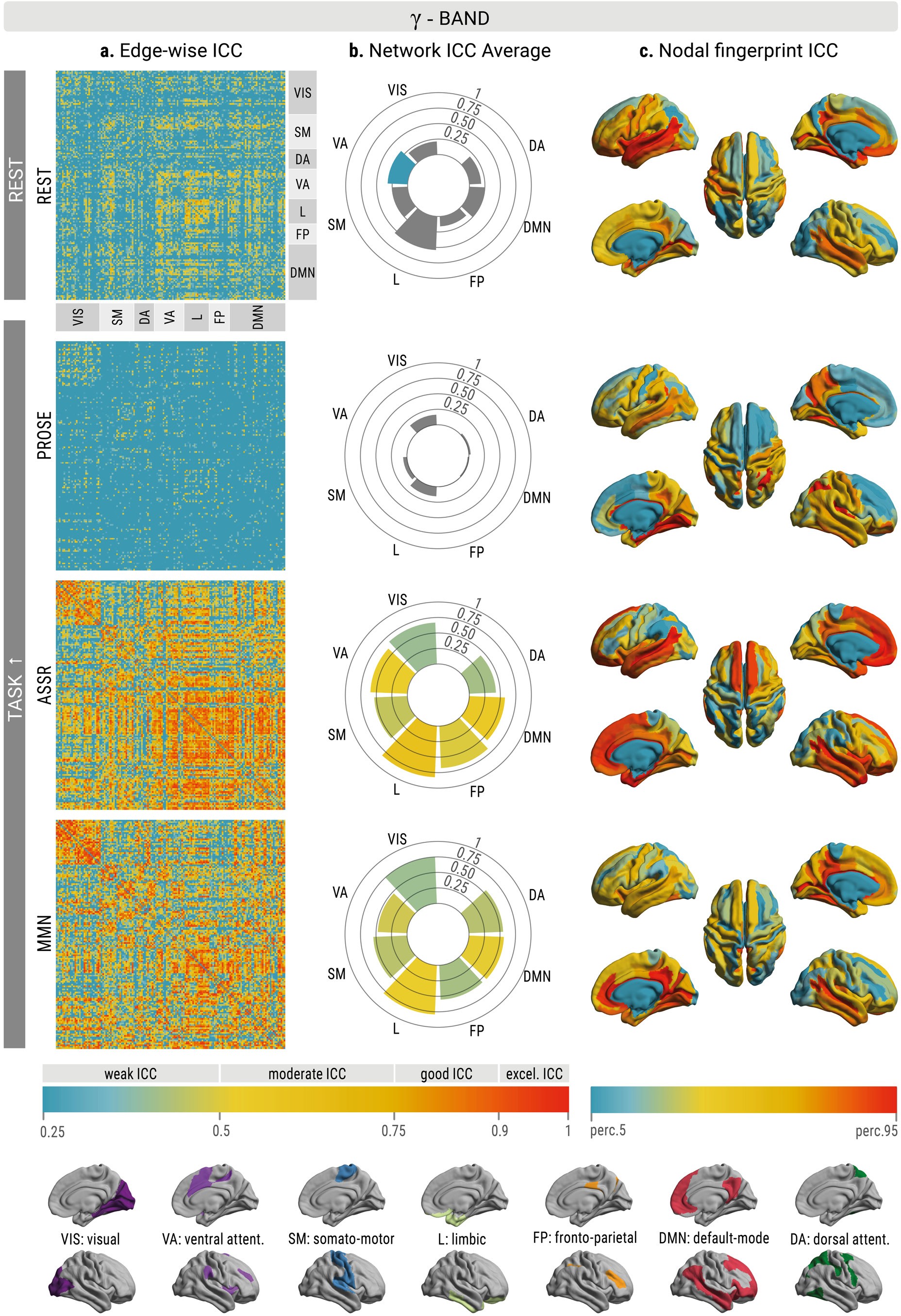

**Supplementary Figure 3 Spatial signatures of individual differentiability in the gamma frequency band.** Edgewise individual differentiability as measured by intraclass correlation (ICC) for each brain state for the gamma frequency band. The ICC values for each of the functional connections per brain state are shown. The higher the value, the more the connection is able to separate an individual from others in a cohort. The brain regions are ordered according to the seven intrinsic functional system organization proposed by Yeo and colleagues. (b) The edgewise ICC scores are averaged within (axis) and between (color) all functional systems of Yeo to better visualize fingerprint patterns within and between functional systems across brain states. Gray indicates an ICC-value < 0.25. (c) Nodal representations of the brain regions involved in individual differentiability during a specific brain state, represented at the 5-95th percentile threshold. The nodal strength of the ICC matrix (i.e., taking the column-wise mean) was used to characterize how central each brain region is for individual differentiation. *Abbreviations of Yeo’s functional networks* VIS = visual; SM = sensorimotor; DA=dorsal attention; VA=ventral attention; L= limbic; FP= frontoparietal; DMN= default mode network.

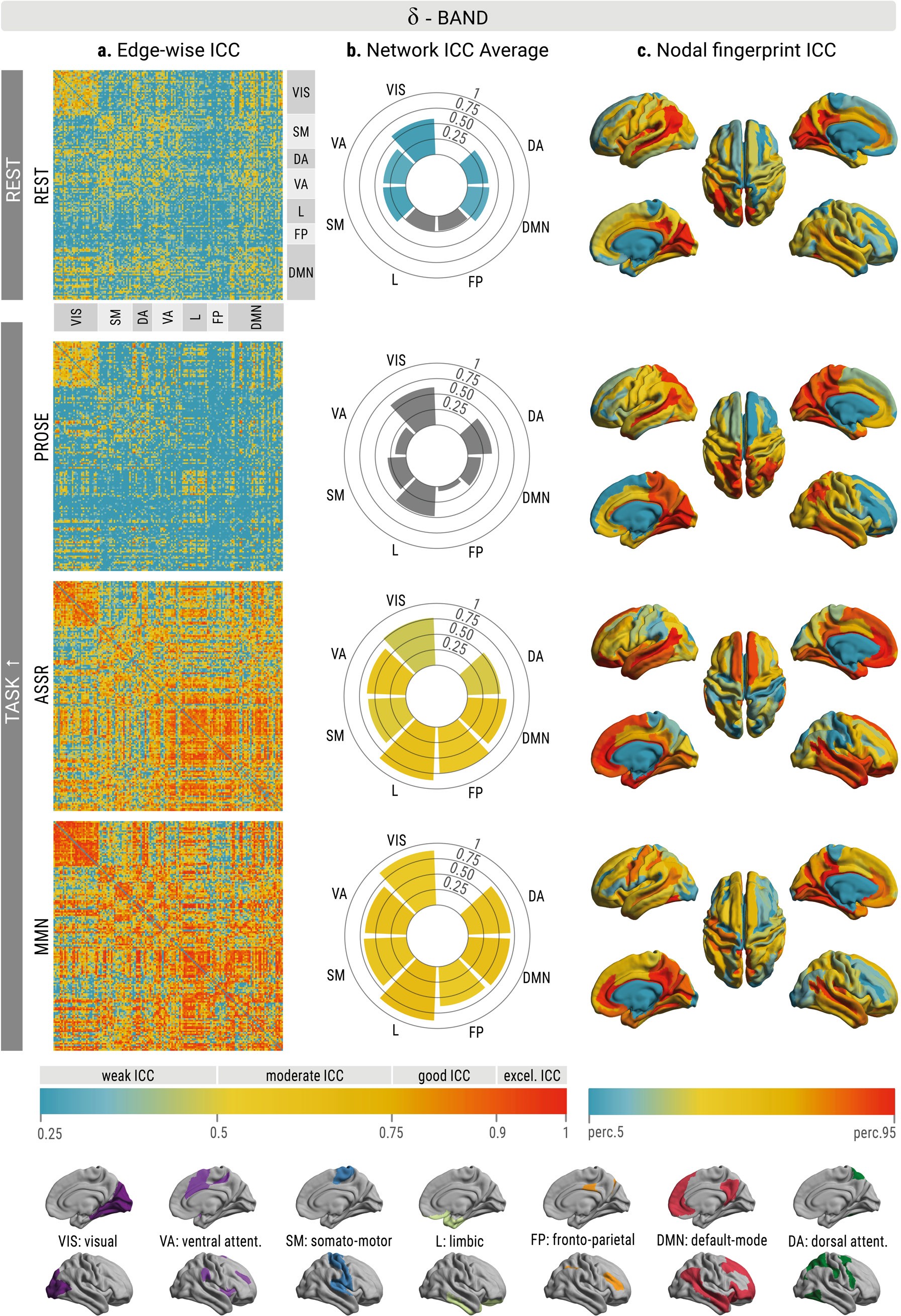

**Supplementary Figure 4 Spatial signatures of individual differentiability in the delta frequency band.** (a) Edgewise individual differentiability as measured by intraclass correlation (ICC) for each brain state for the delta frequency band. The ICC values for each

of the functional connections per brain state are shown. The higher the value, the more the connection is able to separate an individual from others in a cohort. The brain regions are ordered according to the seven intrinsic functional system organization proposed by Yeo and colleagues. (b) The edgewise ICC scores are averaged within (axis) and between (color) all functional systems of Yeo to better visualize fingerprint patterns within and between functional systems across brain states. Gray indicates an ICC-value < 0.25. (c) Nodal representations of the brain regions involved in individual differentiability during a specific brain state, represented at the 5-95th percentile threshold. The nodal strength of the ICC matrix (i.e., taking the column-wise mean) was used to characterize how central each brain region is for individual differentiation. *Abbreviations of Yeo’s functional networks* VIS = visual; SM = sensorimotor; DA=dorsal attention; VA=ventral attention; L= limbic; FP= frontoparietal; DMN= default mode network.

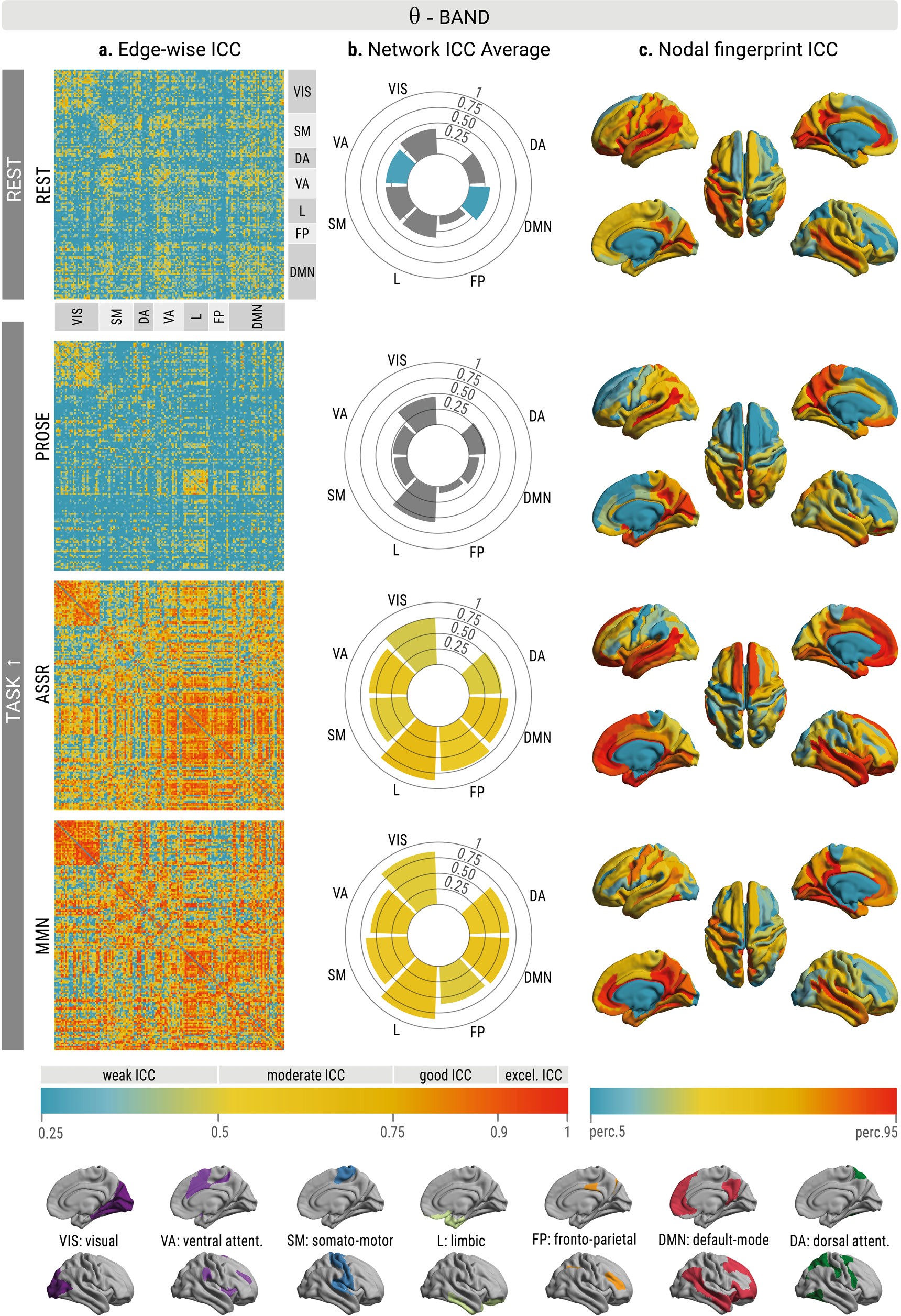

**Supplementary Figure 5 Spatial signatures of individual differentiability in the theta frequency band.** Edgewise individual differentiability as measured by intraclass correlation (ICC) for each brain state for the theta frequency band. The ICC values for each of the functional connections per brain state are shown. The higher the value, the more the

connection is able to separate an individual from others in a cohort. The brain regions are

ordered according to the seven intrinsic functional system organization proposed by Yeo and colleagues. (b) The edgewise ICC scores are averaged within (axis) and between (color) all functional systems of Yeo to better visualize fingerprint patterns within and between functional systems across brain states. Gray indicates an ICC-value < 0.25. (c) Nodal representations of the brain regions involved in individual differentiability during a specific brain state, represented at the 5-95th percentile threshold. The nodal strength of the ICC matrix (i.e., taking the column-wise mean) was used to characterize how central each brain region is for individual differentiation. *Abbreviations of Yeo’s functional networks* VIS = visual; SM = sensorimotor; DA=dorsal attention; VA=ventral attention; L= limbic; FP= frontoparietal; DMN= default mode network.
